# Differential modulation of movement speed with state-dependent deep brain stimulation in Parkinson’s disease

**DOI:** 10.1101/2025.03.19.642627

**Authors:** Alessia Cavallo, Richard M. Köhler, Johannes L. Busch, Jeroen G. V. Habets, Timon Merk, Patricia Zvarova, Jojo Vanhoecke, Thomas S. Binns, Bassam Al-Fatly, Ana Luisa de Almeida Marcelino, Natasha Darcy, Gerd-Helge Schneider, Patricia Krause, Andreas Horn, Katharina Faust, Damian M. Herz, Eric Yttri, Hayriye Cagnan, Andrea A. Kühn, Wolf-Julian Neumann

## Abstract

Subthalamic deep brain stimulation (STN-DBS) provides unprecedented spatiotemporal precision for the treatment of Parkinson’s disease (PD), allowing for direct real-time state-specific adjustments. Inspired by findings from optogenetic stimulation in mice, we hypothesized that STN-DBS effects on movement speed depend on ongoing movement kinematics that patients exhibit during stimulation. To investigate this hypothesis, we implemented a motor state-dependent closed-loop neurostimulation algorithm, adapting DBS burst delivery to ongoing movement speed in 24 PD patients. We found a stronger anti-bradykinetic effect, raising movement speed to the level of healthy controls, when STN-DBS was applied during fast but not slow movements, while only stimulating 5% of overall movement time. To study underlying brain circuits and neurophysiological mechanisms, we investigated the behavioral effects with MRI connectomics and motor cortex electrocorticography. Finally, we demonstrate that machine learning-based brain signal decoding can be used to predict continuous movement speed for fully embedded state-dependent closed-loop algorithms. Our findings provide novel insights into the state-dependency of invasive neuromodulation, which could inspire advanced state-dependent neurostimulation algorithms for brain disorders.

## Introduction

Subthalamic deep brain stimulation (STN-DBS) is an effective treatment for Parkinson’s disease (PD), one of the fastest-growing neurodegenerative disorders^1,2^. While stimulation is traditionally applied in a chronic manner, STN-DBS holds the potential to be adapted within milliseconds, allowing neuromodulation with unprecedented temporal precision^3^. This precision may be utilized to selectively modulate certain brain states, potentially increasing treatment efficacy beyond chronic STN-DBS. In the last decade, subthalamic beta-band activity has been identified as a biomarker of symptom severity^4–6^, leading to the development of beta-adaptive control strategies, which tie stimulation strength to concurrent symptom severity^7–9^. In the context of adaptive DBS, however, it is unknown how concurrent behavior, such as voluntary movement, during adaptive stimulation episodes interacts with the stimulation and potentially modulates its effect. Recent studies in rodents have revealed that transient basal ganglia pathway modulation influences future behavior in a state-specific manner^10,11^. Employing closed-loop speed-triggered optogenetic stimulation of direct spiny projection neurons in the dorsal striatum, Yttri & Dudman (2016) demonstrated that the speed during stimulation was decisive for the behavioral after-effect^10^. Namely, stimulation during the fastest movements led to a subsequent increase in movement speed while stimulation during the slowest movements led to a decrease in speed. A similar reinforcement of movement speed was reported by Markowitz et al. (2023) who additionally showed that both the speed of movements and their occurrence could be enhanced by movement-selective optogenetic stimulation of dopamine-releasing neurons targeting the dorsal striatum. Although these studies target different neuronal populations, both increase the strength of the direct basal ganglia pathway, shifting the balance away from the indirect pathway. STN-DBS applied in Parkinson’s disease is currently thought to modulate indirect pathway activity and net inhibitory output through suppression of hyperdirect input and local synaptic depression^12,13^. On a circuit level, transient STN-DBS may thus have similar effects to transient optogenetic activation of striatal direct pathway or dopaminergic neurons^14^ and could lead to a phasic reduction of pallidothalamic inhibition. With this in mind, we attempt to reproduce the effects of state-specific optogenetic stimulation in human patients using STN-DBS. To do so, we translated the experimental approach presented by Yttri & Dudman (2016) into a motor state-dependent speed-selective STN-DBS paradigm for Parkinson’s disease. Importantly, slowness of movement, also termed bradykinesia, is a cardinal symptom of PD^15,16^. We hypothesized that state-dependent stimulation may have stronger anti-bradykinetic effects when applied during fast movements compared to slow movements. Indeed, stimulation of fast but not slow movements successfully counteracted bradykinesia, providing initial evidence that STN-DBS motor effects are acutely motor state-dependent. Using MRI-based connectomics we identified critical brain networks associated with the speed-specific stimulation effects, revealing supplementary motor cortex and putamen as important hubs. Finally, we uncovered stimulation-induced oscillatory changes in cortical electrocorticography signals and demonstrated their utility in movement decoding for potential future applications of movement-responsive closed-loop STN-DBS.

## Results

### Speed-selective subthalamic deep brain stimulation

To investigate the effects of speed-selective adaptive stimulation we recruited 24 patients with Parkinson‘s disease (PD) undergoing subthalamic deep brain stimulation (STN-DBS). Experiments were performed in the perioperative period before implantation of the pulse generator, which allowed closed-loop control of STN-DBS amplitude through externalized leads. Patients were tested in the clinical OFF state, after 12 h of withdrawal of all dopaminergic medication. Before the start of the paradigm, clinically effective stimulation contacts and amplitudes were determined in a monopolar review. Additionally, 14 healthy age-matched control subjects were recruited to provide a behavioral reference. All subjects were instructed to perform a visuomotor tablet task on a hybrid monitor and digitizing tablet. In this task, a colored rectangle alternately appeared on the left and right side of a tablet screen and subjects performed a forearm movement to guide a touchscreen pen to the target rectangle (Figure 1A). Speed-selective DBS targeting fast or slow movements was applied during a block of 96 movements, followed by a recovery block of the same length during which no stimulation was applied (Figure 1B). The order of conditions was alternated between subjects, who were blinded to the stimulation condition. Healthy participants performed the task without stimulation. The paradigm approximately translated the experimental approach of speed-selective optogenetic stimulation in rodents^10^, to investigate the behavioral after-effects of speed-selective STN-DBS in PD patients. For that, real-time movement speed tracking was used to deliver speed-selective stimulation in PD patients, eliciting 300 ms bursts of bilateral 130 Hz STN-DBS at clinically effective amplitudes (Table S1) during relatively fast or slow movements. The stimulation control algorithm was optimized to stay unbiased by naturally occurring fluctuations in the patient’s motor behavior such as the bradykinetic decrement and at the same time approximated the same portion of stimulated movements in the optogenetic experiment (one-third). In brief, fast and slow movements were classified based on their peak speed relative to the previous two movements. If the peak speed of the ongoing movement exceeded the speed of the previous two movements, it was classified as fast, if it fell under the previous two peak speeds, it was classified as slow (Figure 1A). This dynamic classification algorithm had an accuracy of 96 % across conditions and patients (Table S2) and resulted in the stimulation of 26.3 ± 3.2 % of movements in the fast stimulation block and 32.8 ± 4.5 % of movements during the slow stimulation block across patients. Post hoc analysis revealed that across patients the average peak speed of the stimulated slow and fast movements lay in the 21-37 % and 66-80 % percentile of all movements in the stimulation block, respectively (Figure 1C). Thus, the dynamic classification of fast and slow movements successfully targeted the upper and lower third of the distribution per stimulation block, respectively. Given the short duration of the stimulation bursts, this translated into an active stimulation time of only 8.51 ± 1.46 seconds across the stimulation blocks, roughly 5 % of the block duration (Figure 1D).

**Figure 1:**
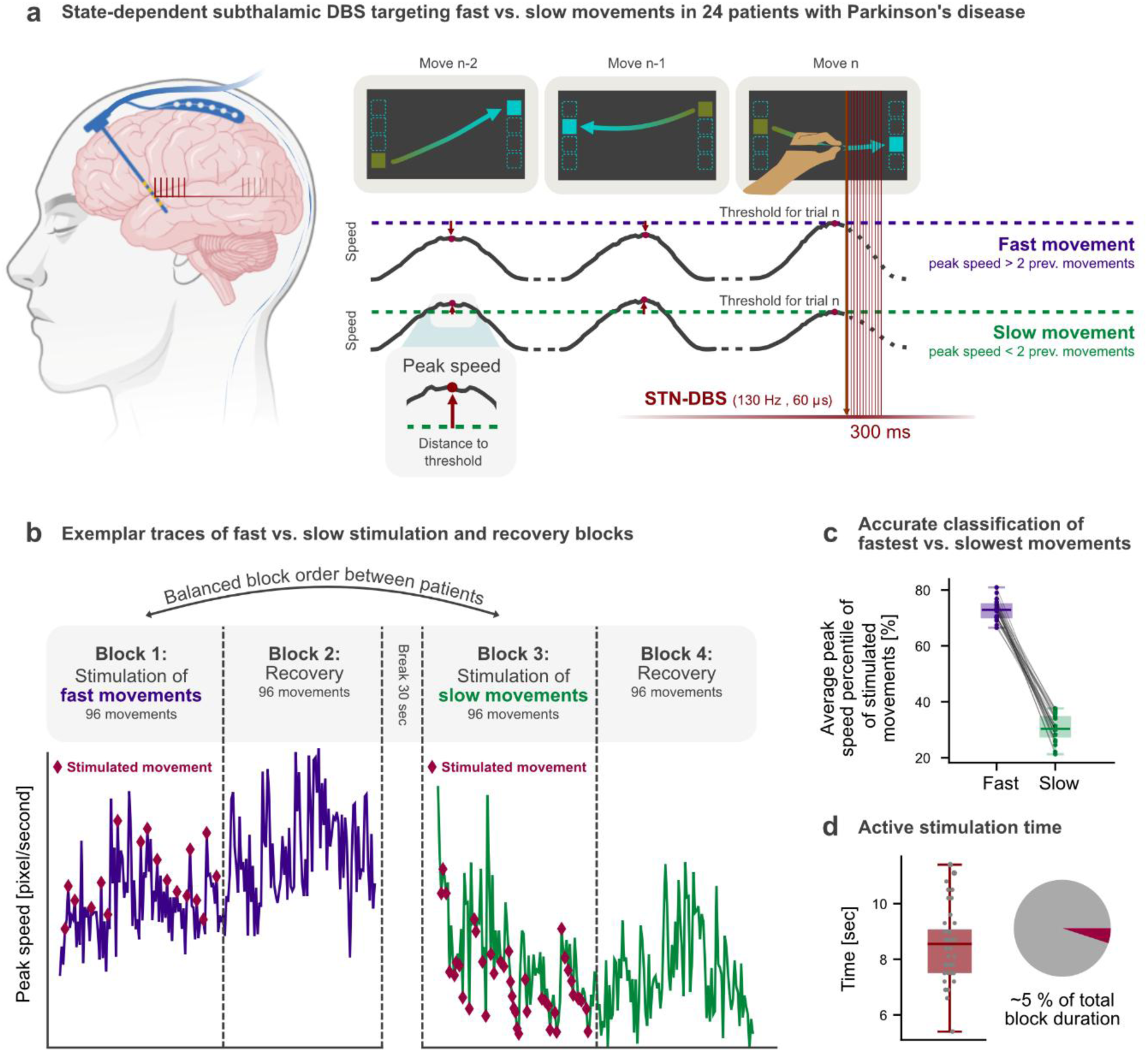
Overview of behavioral task and stimulation algorithm. Participants engaged in a visuomotor tablet task (A) during which they had to move a pen to a colored rectangle appearing on either side of the tablet. In Parkinson’s disease patients OFF dopaminergic medication, 300 ms bursts of subthalamic DBS were applied bilaterally either during movements classified as fast or slow. Stimulation was triggered in accordance with the peak speed, which was used for the classification of fast (purple) and slow (green) conditions. In case of a peak speed higher than that of the previous two movements, the ongoing movement was classified as fast while a peak speed lower than that of the previous two movements led to a classification as slow. (B) Patients underwent 4 blocks of 96 movements each. During blocks 1 and 3 either slow or fast movements were stimulated (red diamonds mark stimulated movements for one exemplary patient), followed by recovery blocks 2 and 4 during which no stimulation was administered. The order of stimulation conditions was balanced between patients. (C) The average peak speed of the stimulated slow and fast movements lay in the 21-37 % and 66-80 % percentile of all movements in the stimulation block, respectively. (D) Average total stimulation time of 8.51 ± 1.46 seconds resulted in stimulation being present during approximately 5 % of the stimulation block.

### Movement state-dependent STN-DBS differentially modulates movement speed

First, we aimed to address the hypothesis that stimulation effects on motor output depend on the ongoing motor behavior, rendering STN-DBS acutely motor state-dependent. Bradykinesia, or slowness of movement, is a disease-defining feature of Parkinson’s disease, leading to a significant deterioration in the amplitude and velocity of movements with repetition. If our hypothesis held true, stimulating only the fastest movements could reinforce high movement speeds and thus counteract bradykinesia, while stimulating particularly slow movements may have no or even an aggravating effect on movement slowness. To quantify the modulation of speed, we extracted the average speed for each movement and calculated the change in percent with respect to the five initial movements of each stimulation block. For healthy control subjects, since no stimulation was applied, the change in speed was averaged over blocks. Indeed, we found that short bursts of STN-DBS during fast compared to slow movements had a stronger anti-bradykinetic effect on overall movement speed i.e. speed declined less strongly in the block in which fast movements were targeted compared to the block in which slow movements were stimulated (Figure 2A; p < 0.01). Comparison with healthy age-matched control subjects revealed that the changes in speed between PD patients in the fast stimulation block and healthy subjects were indistinguishable (Figure 2B; p > 0.05). If slow movements were stimulated, however, the speed of PD patients declined more strongly than that of healthy controls (Figure 2B; p < 0.01). This is the first demonstration that STN-DBS effects on motor control depend on the exhibited movement kinematics at the time stimulation is applied. To test whether the observed effects outlasted the stimulation block, we conducted the same comparisons for the recovery block, during which no stimulation was applied. The difference between stimulation conditions remained significant (Figure 2A; p < 0.05), with a more positive change in speed in the recovery block following the stimulation of fast movements compared to the stimulation of slow movements. No difference was found between the change in speed of healthy control subjects and that of PD patients in the recovery block following either condition (Figure 2B; p > 0.05).

**Figure 2:**
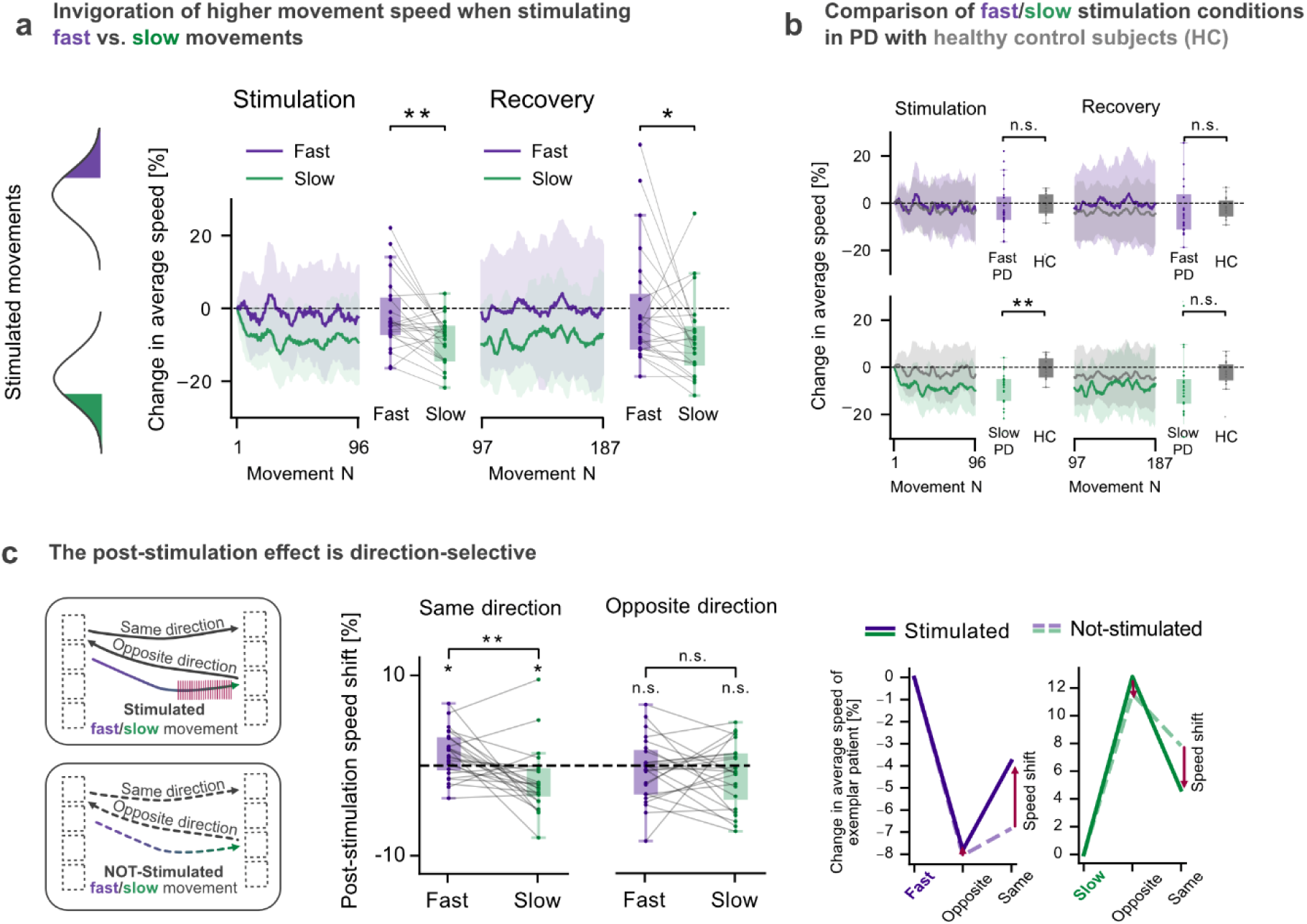
The effect of subthalamic DBS on future movement speed depends on the speed of the stimulated movement. (A) 300 ms STN-DBS had stronger anti-bradykinetic effects with less decline in overall movement speed if applied during fast compared to slow movements. This less pronounced speed decline could also be observed in the recovery block, following the same pattern even though no stimulation was applied. (B) Comparison with age-matched healthy control subjects showed no significant difference in speed changes between PD patients in the fast stimulation block and the control group. However, when slow movements were stimulated, PD patients experienced a significantly greater decline in speed compared to the controls (C). Inspection of post-stimulation movements revealed a stimulation-induced shift towards the speed of the stimulated movement only when subsequent movement direction was concordant with the direction during stimulation. Compared to not-stimulated fast/slow movements extracted from the recovery block stimulation of fast movements led to an increase while stimulation of slow movements resulted in a decrease in subsequent speed. n.s. not significant, * p < 0.05, ** p < 0.01

Our results demonstrate that motor state-dependent STN-DBS differentially modulates movement speed. Stimulation during fast but not slow movements had a significant anti-bradykinetic effect despite stimulation being active for only ∼5% of overall movement time. Notably, stimulation-induced effects outlasted the stimulation block, suggesting plasticity-like mechanisms, which could be exploited in future applications of closed-loop deep brain stimulation for the treatment of PD and other brain disorders.

### Differential speed modulation effects are reach-direction selective

After analyzing block-averaged changes in movement speed, we aimed to characterize the effect of stimulation on individually identified subsequent movements, both in the same and the opposite direction. To do so, we calculated the change in speed for the two following movements normalized to the speed of the stimulated movement. To reveal stimulation-induced shifts in movement speed, we extracted fast and slow movements from the recovery block, during which no stimulation was applied, according to the same algorithm used for speed-selective stimulation. We then calculated the post-stimulation speed shift as the difference between the change in speed following stimulated and not-stimulated movements. We found that stimulation during fast movements resulted in increased movement speed compared to stimulation of slow movements (Figure 2C; p < 0.01). One-sample tests confirmed the expected differential effects, showing significantly increased speed following stimulation of fast movements (p < 0.05), and significantly decreased speed following stimulation of slow movements (p < 0.05). This differential modulation was limited to the subsequent movement in the same direction and was not apparent for the movement in the opposite direction (p > 0.05).

These results indicate that the observed effects are selective for the reach direction at the time point of stimulation, suggesting that state-specific reinforcing effects on motor performance may be movement trajectory-specific.

### MRI connectivity to supplementary motor area and putamen account for acute motor state-specific DBS effects

After examining the behavioral effects of stimulation, we aimed to identify critical brain networks associated with the state-dependent modulation of STN-DBS on movement speed. We used fMRI-based connectomics to elucidate whether the extent of the effect was mediated by the circuit architecture, connecting the stimulated tissue in the STN to the rest of the brain. To do so, we followed the analysis pipeline provided by Lead-DBS, which has been utilized in the past to explain variance in clinical and behavioral outcomes following DBS^17–21^. First, we localized DBS electrodes and determined the volume of activated tissue in each hemisphere for 23 patients based on established methodology (Figure 3A; one subject had an incompatible electrode type)^22^. We then used an openly available connectome derived from resting-state functional MRI images from Parkinson’s disease patients to calculate patient-individual connectivity maps seeding from the stimulation volumes. For each patient, we determined the extent of the state-dependent stimulation effect as the difference in block-averaged speed modulation between fast vs. slow stimulation conditions. For each voxel in the brain, we then correlated the patient-individual effect strengths with the individual functional connectivity, yielding a whole-brain correlation map (Figure 3B). This map constitutes the ‘optimal’ functional connectivity profile associated with the strongest stimulation effect. The similarity of patient-specific connectivity maps with this ‘optimal’ map could significantly account for speed-dependent stimulation effects in left-out patients (Figure 3B, R² = 0.3, p = 0.007, leave-one-patient-out cross-validation). Thus, the stronger an electrode was connected to the identified network, the stronger the state-dependent stimulation was in our experiment. To gather further insights on the key hubs underlying this correlation, we conducted a region-of-interest analysis focused on the human motor network, for which we selected 12 brain regions comprising the cortical and subcortical motor system, including motor cortex, thalamic, basal ganglia, and cerebellar connections^23–26^. For each region, a linear regression model predicting the stimulation effect was trained using the average functional connectivity seeding at the stimulation volumes. Functional connectivity to the supplementary motor area (SMA) best accounted for speed-dependent differences in DBS effects, explaining 36% of the observed variance (Figure 3C; R² = 0.36, p < 0.05, FDR corrected), followed by basal ganglia (putamen, substantia nigra, globus pallidus externus and internus), ventrolateral anterior nucleus of thalamus and dentate nucleus of cerebellum (all p < 0.05, FDR corrected).

**Figure 3:**
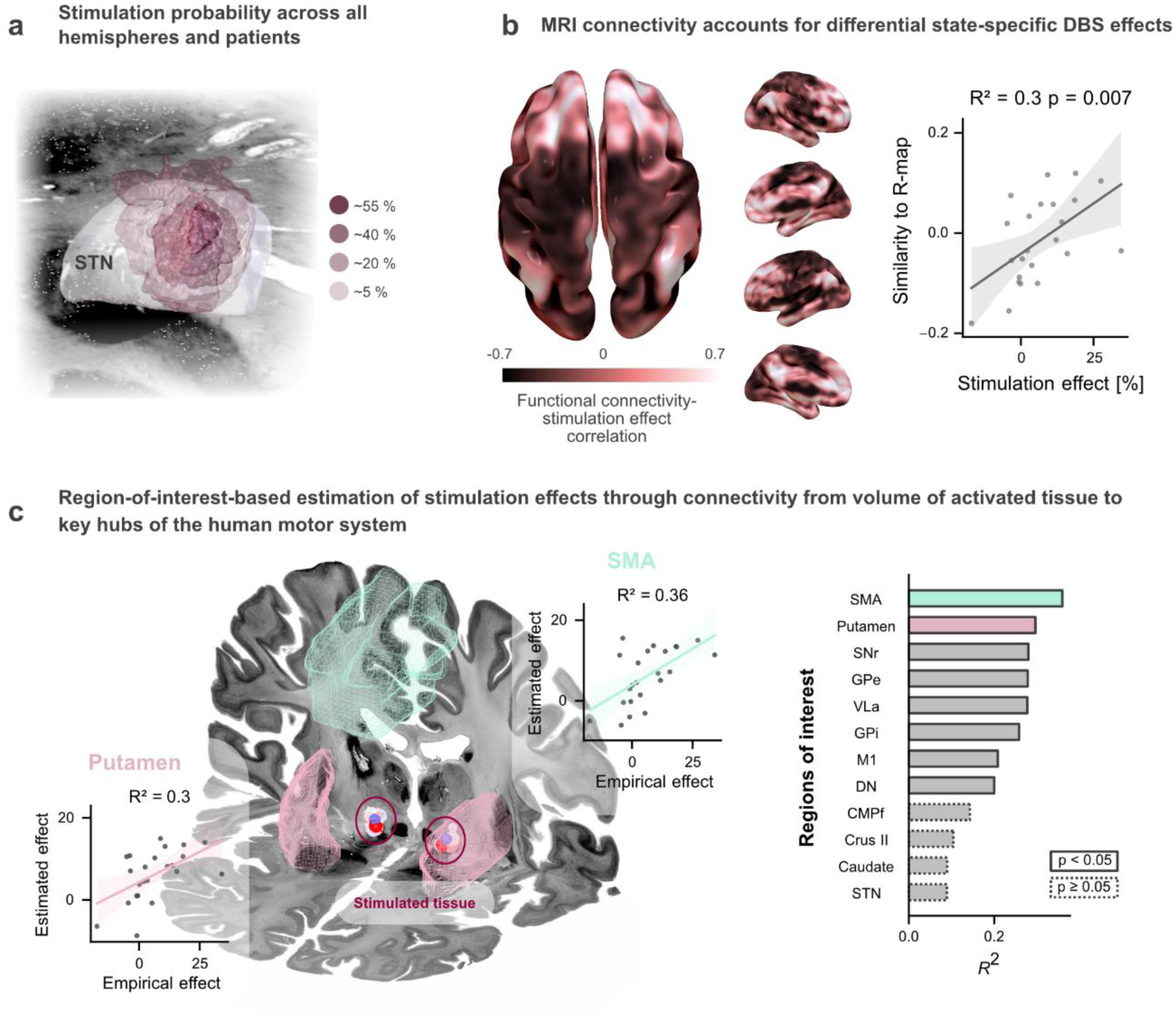
DBS connectivity accounts for speed-dependent stimulation effects. (A) Stimulation probability averaged across hemispheres and patients derived from an overlap of stimulation volumes indicates the highest stimulation probability in the dorsolateral motor part of the STN. (B) The whole-brain voxel-wise correlation map demonstrates the optimal functional connectivity profile for speed-selective DBS effects on movement speed. The more the individual whole-brain functional connectivity profile seeding at the bilateral stimulation volumes matched the ‘optimal’ map, the higher the difference between stimulation conditions (R² = 0.3, p = 0.007, leave-one-subject-out cross-validated). (C) In an additional region-of-interest-based analysis, functional connectivity to the supplementary motor area (SMA) (linear regressor, R² = 0.36) could best predict the observed speed-dependent effect followed by connectivity to the putamen (linear regressor, R² = 0.3) and other basal ganglia nuclei, thalamus, M1 and cerebellar dentate nucleus, which provided a less predictive but still significant estimations (separate linear regression models, solid line indicates a significant regressor, dotted lines indicate non-significance, p-values FDR corrected).SNr: substantia nigra pars reticulata, GPe: globus pallidus externus, VLa: basal ganglia receiving ventrolateral anterior thalamic nucleus, GPi: globus pallidus internus, M1: primary motor cortex, DN: cerebellar dentate nucleus, CMPf: intralaminar centromedian/parafascicular thalamic nucleus, STN: subthalamic nucleus, Crus II: Cerebellar crus II

These results suggest that our speed-selective neuromodulation effect critically depends on the stimulation electrode being connected to a specific brain network, which includes the supplementary motor area, basal ganglia and basal ganglia receiving ventrolateral anterior thalamic nucleus (VLa).

### State-specific neurostimulation modulates motor cortical beta oscillations

To study the neurophysiological underpinnings of the observed stimulation effects, we analyzed electrocorticography (ECoG) signals from an ECoG strip electrode (AdTech) over the right sensorimotor cortex of a single patient undergoing the visuomotor task with their left arm (Figure 4A). Time-resolved beta power between 20-35 Hz was extracted with Morlet wavelets from the channel over the motor cortex located closest to the central sulcus. We then compared the peri- and post-movement changes in oscillatory power between stimulated and not-stimulated movements (Figure 4B). All trials were aligned to the same movement period with matching kinematics. Artifacts at the edges of the DBS burst resulting from switching DBS on and off were excluded. Comparison of stimulated to non-stimulated trial activity revealed a significant reduction in beta power in the peri-movement window (0.1 – 0.3 s; p < 0.05). In contrast, analysis of the post-movement window (0.4 – 1 s) showed a significant stimulation-induced increase in beta power (p < 0.05), after the cessation of stimulation. These findings demonstrate that subthalamic DBS bursts as short as 300 ms can 1) instantaneously suppress beta oscillations in the motor cortex during movement and 2) enhance post-movement beta rebound, previously suggested to relate to cortical reorganization and acute motor adaptation through learning^27–29^. Notably, neither peri-movement beta suppression nor post-movement beta enhancement was pro- or anti-kinetic per se, thus associated with changes in instantaneous kinematics (Figure S1). Instead, we interpret these findings as neurophysiological correlates of state-specific reinforcement of ongoing motor states.

**Figure 4:**
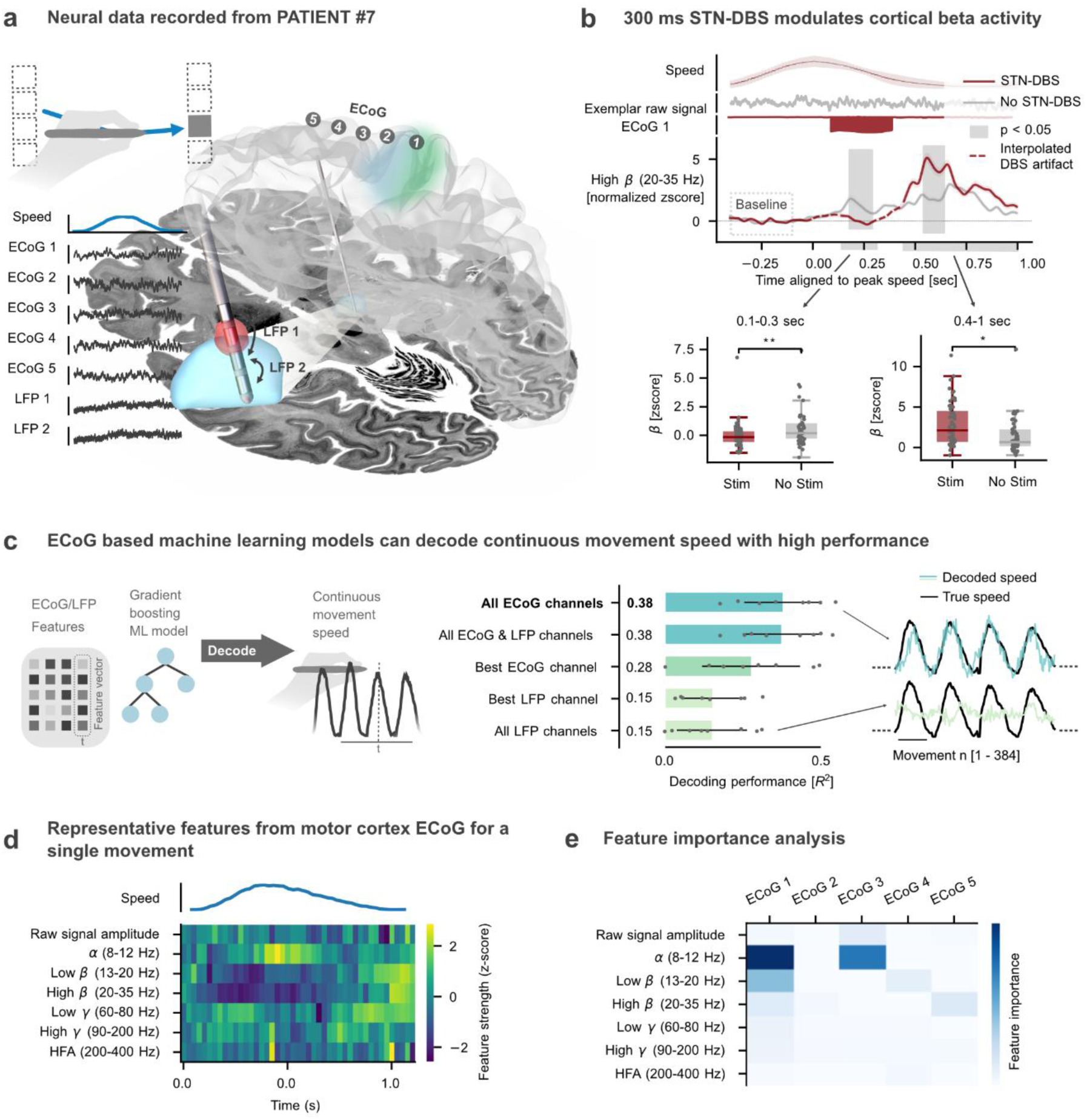
Motor cortex electrocorticography from patient #7. (A) Electrocorticography (ECoG) and DBS/Local field potential (LFP) lead placement in the right hemisphere of patient #7, who performed the motor task with the left hand. The red marker indicates the stimulation contact used for 300 ms bursts of speed-selective stimulation at an amplitude of 2 mA using 130 Hz frequency and 60 µs pulse-width. Common-average ECoG and bipolar LFP referencing resulted in 7 available intracranial electroencephalography (EEG) channels. (B) Comparison of motor cortex beta activity (ECoG channel 1) during stimulated and not-stimulated movements demonstrates a stimulation-induced decrease in beta power during stimulated movements and a stimulation-induced increase in beta power after stimulated movements. Shaded colored areas show the standard error of the mean. Shaded light gray areas indicate clusters of significant differences. Significant differences across averages of movement (0.1 – 0.3 s) and post-movement (0.4 – 1.0 s) periods (gray bars below x-axis) are shown as box plots with individual beta amplitudes below. (C) Continuously recorded movement speed could be successfully decoded using LFP and ECoG signals in gradient boosting machine learning (ML) model (CatBoost). The highest performance was achieved by combining all ECoG channels (R² = 0.38 ± 0.13, hyperparameters optimized and model validated through 4-fold inner and 8-fold outer nested-cross validation) (D) Exemplar trace of oscillatory features and the raw signal amplitude extracted from an exemplar channel and movement using py**_**neuromodulation. (E) Analysis of ML model feature importance (prediction value change) reveals alpha and low beta power as the most informative features for movement speed decoding (optimal decoder for exemplar outer fold). n.s. not significant, * p < 0.05, ** p < 0.001

### Towards fully embedded state-specific stimulation through movement speed decoding from invasive neurophysiology

Our behavioral finding showing a less pronounced bradykinetic decline from stimulation of fast compared to slow movements highlights the potential of acute motor state-specific DBS approaches for the treatment of PD in the future. The use of external sensors to infer behavioral states, however, such as the cursor position employed in the present study, has limited practicality for chronic treatment of patients. Therefore, we aimed to investigate whether invasively recorded ECoG and local field potential (LFP) signals could be used to decode the motor behavior in the employed tablet task. The signals from five common-average-referenced ECoG and two bipolar LFP channels were used to predict the movement speed of a single patient (maximum number of channels available, Figure 4A). Six oscillatory features [alpha (8–12 Hz), low beta (13–20 Hz), high beta (20–35 Hz), low gamma (60–80 Hz), high gamma (90–200 Hz) and high-frequency activity (HFA) (200-400 Hz) power] and the raw amplitude were extracted from the signal using the real-time compatible pipeline provided by py_neuromodulation^30^ (Figure 4D). Based on these features, a gradient boosting machine learning model (CatBoost^31^) was trained to predict the continuously recorded movement speed of the patient (Figure 4C). Importantly, only features from previous time points were used to inform the prediction of the instantaneous movement speed. The decoding performance of different channels and channel combinations was tested in a large-scale Bayesian optimization hyperparameter search (see Table S3). A combination of all ECoG channels emerged with the highest performance (R² = 0.38), similar to the combination of all available ECoG and LFP channels (R² = 0.38). Notably, the best ECoG channel (R² = 0.28), achieved higher decoding performances than both LFP channels combined (R² = 0.15; Figure 4C, average performance from an 8-fold nested cross-validation). Analysis of feature importance revealed that alpha and low beta power estimates in the ECoG channel, located over the motor cortex, were the most informative for the decoding of movement speed (Figure 4E). These results provide a proof of principle that continuous movement speed can be decoded from motor cortex ECoG signals with high performance, which may pave the way for the development of fully embedded speed-dependent DBS approaches in the future.

## Discussion

Our study implements a novel motor-state-dependent closed-loop neurostimulation approach to provide an important proof of principle, showing that motor effects of STN-DBS depend on the acute motor state during which stimulation is applied. We derive three major findings from our study. First, we demonstrate that STN-DBS applied in brief bursts with closed-loop control does not have a “prokinetic” effect per se. Instead, we found that the same stimulation can evoke differential behavioral effects depending on concurrent behavior. Specifically, we found a more pronounced anti-bradykinetic effect by stimulating fast compared to slow movements, shifting the speed of the subsequent trial towards the speed of the stimulated movement, compatible with a reinforcement effect of closed-loop STN stimulation. Next, using normative functional connectomes we elucidate brain-wide networks associated with this differential modulation, identifying the supplementary motor area and basal ganglia as key hubs for the induction of the state-dependent stimulation effect. Finally, using intracranial cortex electrophysiology in a single patient, we report a stimulation-induced decrease in peri-movement and an increase in post-movement cortical beta activity, suggesting that 300 ms STN-DBS modulates cortical processing associated with motor adaptation. In the same patient, sensorimotor ECoG signals could be used to decode continuous movement speed with high performance, supporting the feasibility of fully embedded movement-dependent DBS approaches. Altogether, we argue that the state during STN-DBS influences the behavioral after-effects of stimulation, with stimulation of more vigorous motor behavior potentially providing a more effective alleviation of bradykinesia than stimulation applied during less vigorous motor states. This has important implications for the development of future DBS control algorithms, motivating a speed-adaptive approach for maximization of therapeutic efficacy.

### Reinforcement of behavior through basal ganglia neuromodulation: From rodents to humans

Traditionally, the basal ganglia have been divided into a prokinetic direct “Go” pathway and an antikinetic indirect “NoGo” pathway. Consequently, high-frequency STN-DBS, suppressing activity in the antikinetic indirect pathway, has been interpreted as being prokinetic per se. Recent work in rodents, however, challenges the dichotomy of solely pro-and antikinetic basal ganglia pathways by showing that both pathways are intrinsically active during movement initiation and that optogenetic stimulation of both pathways can invigorate or weaken movement^10,32^. Importantly, this bidirectional modulation was reported to depend on the concurrent movement state, with stimulation of direct striatal neurons leading to an increase in speed if fast and a decrease in speed if slow movements were stimulated^10^. Studies modulating basal ganglia activity through manipulation of dopamine signaling further underline the state-dependency of basal ganglia processing^11,33^. Here, Markowitz et al. (2023) reported that exogenously induced dopamine transients in the dorsal striatum can reinforce the occurrence of the ongoing behavioral state in the future^11^. Our study reports acute motor state-dependent modulation of motor behavior through STN-DBS in humans, indicating that its effect might go beyond a purely prokinetic mechanism and instead follow the logic of phasic dopaminergic control^14^. Stimulation during fast movements resulted in a more positive modulation of movement speed than stimulation targeting slow movements. Looking more closely, we found that STN-DBS led to reinforcement-like shifts in future movement speed in a direction- and thus potentially neural pathway-specific manner. While our findings are in line with the results from the speed-selective optogenetic stimulation in rodents, key differences merit further elaboration. As our study was performed with human PD patients, we could not optogenetically stimulate striatal neurons but instead applied high-frequency STN-DBS. We argue that high-frequency STN-DBS may have a similar influence on net basal ganglia output as the activation of striatal pathway neurons^14^. Here, optogenetic stimulation of D1-dopamine receptor-expressing neurons can shift the basal ganglia pathway balance away from the indirect pathway by increasing activity in the direct pathway. Similarly, STN-DBS can also modulate the pathway balance in favor of the direct pathway, not through activation of the direct pathway but through suppression of the STN, the main driver of the indirect pathway^12,34,35^. On the behavioral level, despite only stimulating a fraction of the experiment time, we observed lasting changes to movement speed in dependence on ongoing kinematics. How exactly this STN-DBS-induced modulation gives rise to changes in future motor behavior, however, can only be speculated and may involve plasticity at cortico-striatal, thalamo-striatal, thalamo-cortical, and cortico-cortical synapses^14,36,37^. Regardless of the underlying mechanism, our study provides evidence that motor effects of STN-DBS depend on ongoing movement kinematics and that bradykinesia, the cardinal feature of PD may be best treated by stimulating during fast and vigorous motor output.

### Speed-selective STN-DBS engages brain-wide networks with supplementary motor area and basal ganglia as key hubs

Over the past decade, MRI-based DBS connectomics has emerged as a powerful tool to uncover circuit effects of invasive neuromodulation in humans^20,38^. Previous studies have focused on predicting patient-specific clinical or behavioral stimulation effects based on the individual location of chronically active stimulation electrodes^17,20,39^. In our study, we extended this framework to the differential effect of state-specific stimulation algorithms, namely the difference between stimulating fast vs. slow movements. We could identify a whole-brain network related to the stimulation-induced speed-selective effect, elicited by only brief 300 ms bursts of closed-loop DBS. The more electrodes were connected to this network, the stronger their effect was on the identified movement-specific neuromodulation. The network highlighted broad frontal cortical, thalamic and basal ganglia regions, which is in line with previously reported networks mediating clinical DBS effects^17,40^. Beyond whole-brain connectivity, a region-of-interest analysis revealed that SMA and basal ganglia nuclei best accounted for the observed effect. The relevance of these regions is in line with previous studies showing that SMA^41^, as well as basal ganglia^42^, encode movement kinematics such as hand speed and that alterations in activity have been linked to bradykinesia^43^. Moreover, both regions are involved in skill learning^44,45^ and are known to drive changes in future motor behavior. In this regard, it has been hypothesized that “neural reinforcement” in cortico-basal ganglia circuits may refine and strengthen cortical activity patterns associated with successful behavior, ultimately driving skill learning and behavioral consolidation^36^. Our findings can be integrated into this framework, as speed-selective STN-DBS may have reinforced neural patterns associated with either slow or fast movements, potentially associated with cortical beta oscillatory activity as discussed below.

### Speed-selective closed-loop STN-DBS modulates cortical beta power

To further elucidate the remote effects of speed-selective STN-DBS, we investigated the impact of stimulation on motor cortex beta activity. In the STN, excessive beta synchronization has been repeatedly linked to PD symptom severity, which is reduced by DBS^4,46,47^. While evidence focusing on cortical modulations is less consistent, work suggests that STN-DBS attenuates beta power over motor cortices^48^. We extend this observation in a patient who received a contralateral electrocorticography strip, by showing that < 1-second STN-DBS applied during a movement reduces beta power recorded over the motor cortex. Moreover, we report that speed-selective DBS not only influences instantaneous oscillation but also increases post-movement beta synchronization, previously termed beta rebound^49^. While analysis of the present data from a single patient does not allow us to systematically investigate the relationship between cortical oscillatory changes and the strength of speed-dependent DBS effects, we may speculate about the involvement of post-movement beta rebound in the observed state-dependent effect. It has been reported that the strength of beta rebound is linked to the adaptation of motor behavior, with higher levels reflecting maintenance and reduced activity indicating an adaptation of the current motor program^28,50^. Following this rationale, the increase in beta rebound following selective stimulation of fast and slow movement may be associated with the consolidation of ongoing activity patterns and thus increase the likelihood of producing similar behavior, i.e. moving fast after a fast movement has been stimulated. Follow-up studies investigating the relationship between speed-selective DBS effects and oscillatory modulation in a larger patient cohort and animal studies further characterizing the involvement of oscillations in synaptic plasticity in the motor system are needed to elucidate this hypothesis.

### Towards neural reinforcement with state-adaptive deep brain stimulation for Parkinson’s disease and beyond

While chronic DBS has been an effective treatment option for PD for the last 30 years, a growing body of research focuses on the development of dynamically controlled adaptive DBS approaches, increasing treatment efficacy and reducing side effects. Besides more recent work suggesting adaptation based on gamma-oscillations^51,52^, most existing approaches for PD have focused on beta-band activity as a control signal, which has been associated with symptom severity^6^. While the clinical efficacy of beta-adaptive DBS has been reported^53^ and its application is currently being tested in clinical trials, concerns regarding the timing of stimulation have arisen. Subthalamic beta activity is reduced during movement^54,55^, and consequently, beta-adaptive control reduces stimulation amplitudes during movement^56–58^. Our data indicating reinforcement-like effects of STN-DBS motivates an alternative approach that focuses on the motor state as a potential control signal. Recently, Dixon et al. (2024) demonstrated the feasibility and efficacy of movement-adaptive DBS, increasing stimulation amplitude during a movement and decreasing the amplitude during rest periods^59,60^. The idea of triggering DBS during selective behavioral states has also been explored for the treatment of tremor^61,62^. Importantly, these approaches do not rely on external sensors for the detection of motor states but, similarly to beta-adaptive approaches, decode the presence of these states from invasively recorded neural signals. Dixon et al. (2024) for example, classified the occurrence of movement or rest based on the oscillatory power in subthalamic LFP and cortical ECoG channels. Our results encourage the development of adaptive DBS algorithms beyond the binary classification of movement. Here, stimulation amplitude could be controlled based on continuously decoded movement vigor to selectively stimulate and potentially reinforce high-vigor states in PD. It has been shown that grip force strength can be continuously decoded based on subthalamic LFP and cortical ECoG signals^63^. Similarly, we demonstrate in a single patient implanted with ECoG electrodes that the movement speed in the present task can be successfully predicted without the need for external sensors. The fact that we obtained the highest decoding performance using ECoG in addition to STN-LFP signals, aligns with previous work showing the superiority of cortical over subcortical signals for movement decoding^63^. Taken together, our findings motivate the development of speed-adaptive DBS approaches for the treatment of PD and provide a proof of principle that cortical ECoG might be a suitable decoding signal, allowing fully invasive closed-loop control. Beyond PD, reinforcement-like effects of DBS may be leveraged to accelerate learning outside the motor domain. For prosthetics such as brain-spine interfaces^64^, auditory implants^65^, and artificial retinas^66^, state-selective DBS could in the farther future help to strengthen input-output of neural circuits relationships and thus accelerate adaptation to neuroprosthetics.

### Limitations

While the abovementioned implications are promising, it is important to consider the limitations of this study. First of all, limited recording time has hindered us from performing an experimental condition without stimulation. Thus, we cannot report an absolute stimulation-induced increase in movement speed from baseline. Nevertheless, the use of two differential stimulation conditions provides a robust basis to investigate the state-dependent nature of the stimulation effect. The execution in a pseudo-randomized order moreover ruled out the possibility that differences between conditions were influenced by order effects. Second, the strength of this effect varied strongly between individuals, with only a subset of patients showing strongly elevated speed in the fast compared to the slow condition. While a significant amount of this variation could be explained by differences in the stimulated volume, other factors might have attributed to the variability. Electrode implantation is known to induce a ‘stun effect,’ characterized by a temporary suppression of Parkinsonian symptoms, likely resulting from a microlesion in the STN^67^. Different extents of this ‘stun effect’ across patients might have led to different states of the cortex–basal ganglia network and consequently to differential state-dependent DBS-induced processing. Third, only a preliminary clinical review could be carried out to select the stimulation parameters used during the study. Consequently, the strengths of the observed DBS effects may not fully represent the extent potentially obtainable after an extensive review of stimulation parameters. However, even with preliminary parameter optimization, the observed effects indicate a promising direction. Lastly, the results obtained from the ECoG signals in a single patient lack generalizability. Similarly, the discussed mechanisms are merely a speculation and must be interpreted cautiously. However, we argue that the presented results provide a valuable starting point for investigating state-dependent STN-DBS effects.

### Conclusion

In conclusion, our study demonstrates that the motor state during which STN-DBS is applied critically influences its behavioral effects, providing a key proof of principle for state-dependent adaptive stimulation in Parkinson’s disease. We show that closed-loop, motor state-dependent STN-DBS can differentially modulate behavior, with stimulation during faster movements exhibiting a more pronounced anti-bradykinetic effect. This effect seemed maximal when electrodes were strongly connected to the supplementary motor area and basal ganglia, while intracranial electrophysiology in a single patient revealed stimulation-induced changes in cortical beta activity. These findings underscore the potential and feasibility of movement-dependent DBS strategies, suggesting that stimulation applied during more vigorous motor states may enhance therapeutic efficacy. Taken together, our results pave the way for the development of novel speed-adaptive DBS control algorithms, optimizing outcomes for individuals with Parkinson’s disease.

## Acknowledgments

Funded by Deutsche Forschungsgemeinschaft (DFG, German Research Foundation) – Project-ID 424778381 – TRR 295, the Bundesministerium für Bildung und Forschung (BMBF, project FKZ01GQ1802) and the European Union (ERC, ReinforceBG, 101077060). Views and opinions expressed are however those of the author(s) only and do not necessarily reflect those of the European Union or the European Research Council. Neither the European Union nor the granting authority can be held responsible for them.

We thank all patients who participated in this study. Without their dedication to contribute to the understanding and treatment of Parkinson’s disease, our research would not be possible.

## Author contributions

Conceptualization, W.-J.N, A.C.; Methodology, A.C., W.-J.N.; Software, T.M, T.S.B., P.Z.; Data Acquisition: A.C., R.M.K, J.B., J.H.; Formal Analysis, A.C., W.-J.N; Investigation, A.H., D.M.H., E.Y., H.C.: Resources, A.A.K., P.K., K.F., G.-H.S., A.L.A.M, N.D., B.A.; Data Curation, J.V.; Writing – Original Draft, A.C., W.-J.N.; Writing – Review & Editing; A.C., W.-J.N, R.M.K., T.M., J.V., T.S.B., P.Z., J.B., D.M.H., E.Y., H.C., A.L.A.M., N.D., B.A.; Visualisation, A.C., W.-J.N.; Supervision, W.-J.N.; Project Administration, A.A.K., W.-J.N.; Funding Acquisition,W.-J.N.

## Declaration of interests

A.A.K. reports personal fees from Medtronic and Boston Scientific. G.-H.S. reports personal fees from Medtronic, Boston Scientific, and Abbott. W.-J.N. serves as a consultant to InBrain and reports personal fees from Medtronic. A.H. reports lecture fees for Boston Scientific and is a consultant for FxNeuromodulation and Abbott and serves as a co-inventor on a patent application by Charité University Medicine Berlin that covers multisymptom DBS fiberfiltering and an automated DBS parameter suggestion algorithm unrelated to this work. The application has been submitted on July 21, 2023, with the patent office of Luxembourg (application #LU103178).

## Methods

### Participants

Twenty-four patients diagnosed with idiopathic Parkinson’s disease of primary akinetic-rigid motor phenotype with clinical indication for DBS were enrolled at the Department of Neurosurgery at the Charité – Universitätsmedizin Berlin (59.2 ± 9.4 years, 5 female, see Table S1). DBS leads were bilaterally implanted into the STN. In nine patients, a subdural ECoG electrode strip was implanted unilaterally targeting the hand knob area of the primary motor cortex for research purposes. Experimental sessions were conducted in the perioperative state between the first and second surgical intervention and after the withdrawal of dopaminergic medication. Each patient underwent one session, including speed-selective stimulation during a behavioral task and simultaneous electrophysiological recordings. After completion of the experimental paradigm, the patient’s symptom severity was quantified with the Unified Parkinson’s Disease Rating Scale (UPDRS) III (29.0 ± 9.3, see Table S1). Fourteen healthy age-matched participants were additionally recruited to perform the same behavioral task (59.5 ± 6.5 years 9 female, see Table S1).

### Ethics declaration

The study presented in this manuscript was performed according to the standards set by the declaration of Helsinki and after approval by the ethics committee at Charité Universitätsmedizin Berlin (EA2/129/17). All patients and healthy participants provided informed consent to participate in the research. The data were collected, stored, and processed in compliance with the General Data Protection Regulation of the European Union.

### DBS and ECoG placement

DBS implantation followed a two-step approach. In the first surgery, DBS leads were placed stereotactically after co-registration of preoperative MRI and CT images. In nine patients, a single ECoG electrode strip was placed subdurally onto one hemisphere after minimal enlargement (∼2 mm) of the frontal burr hole. The ECoG strip was aimed posteriorly toward the hand knob region of the motor cortex. The ECoG strip was placed ipsilaterally to the implantable pulse generator (right = 8, left = 1). All electrodes were then externalised through the burr holes via dedicated externalisation cables. Patients remained on the ward for a duration of 4 to 7 days until the second surgery. In the second intervention, externalisation cables of the DBS leads were replaced with permanent cables that were tunnelled subcutaneously and connected to a subclavicular implantable pulse generator. ECoG electrodes were removed via the burr hole during the second surgery.

### Anatomical localization of electrodes

DBS and ECoG electrodes were localized using standard settings in Lead-DBS^68^. In brief, pre-operative MRI and post-operative CT images were linearly co-registered using Statistical Parametric Mapping software^69^ (SPM12, https://www.fil.ion.ucl.ac.uk/spm/software/spm12/), corrected for brain shift, and normalized to MNI space (Montreal Neurological Institute; MNI 2009b NLIN ASYM atlas) using default presets of the Advanced Normalization Tools^70^ (ANTs, http://stnava.github.io/ANTs/). DBS electrodes were automatically pre-reconstructed using the PaCER algorithm^71^ for postoperative CT and later manually refined if needed. MNI coordinates of each electrode contact were lastly extracted to be used in spatial analyses of electrophysiological signals.

### Experimental paradigm

Rectangles of 175×175 pixel were displayed on a Wacom Cintiq 16 Tablet (1920×1080 pixel) using Psychtoollbox in MATLAB. Rectangles were placed either at the left end (100 pixel) or the right end of the tablet (1820 pixel) at one of four positions along the y-axis (390, 465, 615, 690 pixel). Patients were instructed to move a pan held in their dominant hand to the displayed rectangle. Upon arrival, the rectangle changed color and disappeared after an inter-movement-interval of 350 ms. At the same time, a rectangle on the opposite side appeared, leading to alternating movements from one side of the tablet to the other. The paradigm consisted of 4 blocks containing 96 movements each. As rectangles were placed at 4 different locations along the y-axis, 32 possible trajectories could be performed, which were pseudo-randomized within each block and kept constant across participants. The cursor position was sampled at 62 Hz using MATLAB and used to calculate movement speed in real time. The difference between consecutive cursor positions was calculated, divided by the elapsed time, and averaged over 6 samples to obtain a smoother read-out. Speed-selective stimulation was applied based on the peak speed of each movement. The peak speed of the trial was extracted in one of two ways: In twenty patients the closest timepoint following the peak speed was detected by three subsequently decreasing speed values and the maximum value of the previous samples was extracted as peak speed. In four patients the time of peak speed after movement initiation was computed as the 80^th^ percentile of 32 additional calibration trials and when reached, the peak speed was extracted for the ongoing movement. The former approach was used for the majority of the patients as it did not require additional calibration trials while maintaining classification accuracy (see Table S2 for group-averaged classification accuracy). To classify movements as slow or fast the peak speed of the ongoing movement was compared to the peak speed of the previous two movements. A movement was classified as fast if the peak speed exceeded that of the previous two movements and it was classified as slow if it fell below the previous two values. In blocks 1 and 3 (stimulation blocks) speed-selective stimulation targeted either slow or fast movements. The order of conditions was balanced across patients. Movements lasted 812 ± 115 ms on average. Stimulation on average started 66 ms after the peak speed and lasted 300 ms. In blocks 2 and 4 (recovery blocks) no stimulation was applied. A 30-second break was introduced between block 2 and block 3 for all patients and an additional 30-second break between blocks 1 and 2 as well as 3 and 4 was present for four out of twenty-four patients. Again, the former approach was used for the majority of the patients to reduce overall recording time.

### Subthalamic deep brain stimulation

Two different DBS lead models with either 8 (n = 23) or 16 (n = 1) contacts were implanted. Three segmented contacts on one level were used for monopolar stimulation. During a unilateral test stimulation preceding the stimulation paradigm, the amplitude was progressively increased, and side effects as well as improvements in rigidity and movement speed of the UPDRS III finger tapping test were visually evaluated. All contacts apart from the top and bottom ones were tested. The contacts on each hemisphere with maximum stimulation amplitude and clinical improvement in the absence of side effects were selected for the subsequent experiment. Side effects during bilateral stimulation were additionally evaluated and stimulation amplitude reduced if needed. Stimulation was applied bilaterally without ramping at 130 Hz with a pulse width of 60 µs and a mean amplitude of 2.4 ± 0.4 (right) and 2.4 ± 0.5 (left) mA (see Table S1 for details).

### Analysis of behavioral data

To extract kinematic measures specific to individual movements, the onset and offset of each movement had to be determined. Centering the data on the time point of peak speed, movement onset and offset were set as the last samples above a threshold of 800.77 pixel/second. The threshold was defined as three standard deviations above the speed during the last 300 ms of the inter-trial rest period aggregated over trials and patients. The average movement speed between movement onset and offset was calculated as the main outcome measure for each trial.

For the analysis of speed modulation in a whole block, the first five trials were excluded from analysis and trials whose average speed exceeded 2413.35 pixels/second were classified as outliers and replaced with NaN values. The outlier threshold was defined as three standard deviations above the average speed aggregated over trials and patients. In order to compare the change in speed between conditions, movement speed was first normalized with respect to the average value of the first five movements by subtracting and subsequently dividing by the average value, yielding the change in speed from block start in %. Speed values in the recovery blocks were normalized to the start of the preceding stimulation block. For statistical comparison, speed values were averaged over trials for each block. For healthy control subjects, speed values were averaged across both stimulation (1 and 3) and both recovery (2 and 4) blocks as no stimulation was applied. Stimulation conditions were statistically compared in a within-subject design using a two-sided paired-sample non-parametric permutation test with 100,000 permutations at an alpha level of 0.05. The average change in speed between healthy control subjects and both stimulation conditions was statistically compared using a two-sided independent-sample non-parametric permutation test with 100,000 permutations at an alpha level of 0.05.

In order to analyze the effect of speed-selective stimulation on subsequent trials, we identified the stimulated and two subsequent movements for each subject. We then normalized the speed in the subsequent trials with respect to the stimulated trial yielding a change of speed in %. As stimulated movements were either fast or slow, subsequent movement speed was shifted in the opposite direction, with speed decreasing after a fast and increasing after a slow movement. To compare stimulation effects independent of this inherent shift, we identified fast and slow movements in the recovery block, during which no stimulation was applied. Exactly as for stimulated movements, the change in speed of the two following movements was calculated. We then subtracted the change in speed following not-stimulated movements from the change in speed following stimulated movements. The difference in the stimulation-induced speed shift between conditions was statistically compared using a two-sided paired-sample non-parametric permutation test with 100,000 permutations at an alpha level of 0.05. To examine whether the speed shift for each condition significantly differed from 0, a sign-flipping permutation test with 100,000 permutations at an alpha level of 0.05 was performed.

### fMRI-based functional connectivity analysis

An openly available Parkinson’s disease fMRI group connectome^17^ (www.lead-dbs.org) derived from the Parkinson’s progression markers initiative (PPMI) database was used to investigate the relationship between speed-dependent stimulation effects and fMRI resting-state connectivity on the whole-brain level. All scanning parameters are published on the website (www.ppmi-info.org). Stimulation volumes were computed using the Simbio/Fieldtrip model in the Lead-DBS toolbox for 23 out of 24 patients (Patient 4 was excluded because the implanted DBS lead model was not supported). Using the bilateral stimulation volumes as seeds, whole-brain functional connectivity maps were generated for each patient. Then, the connectivity values and the stimulation effects were correlated across patients for each voxel. This resulted in an R-map representing the whole-brain functional connectivity profile associated with the strongest stimulation effect. The stimulation effect was quantified as the difference between the average change in speed in % during blocks 1 & 2 (stimulation of fast movements and subsequent recovery) and the average change in speed during blocks 3 & 4 (stimulation of slow movements and subsequent recovery). The explanatory power of the R-map was validated using a leave-one-patient-out approach: For each patient, a distinct R-map was calculated based on the maps of the remaining patients which was then spatially correlated with the connectivity map of the left-out patient. The resulting network score, describing the similarity of the patient’s connectivity profile to the optimal map, was then correlated with the observed stimulation effects. To investigate how individual brain regions contribute to explaining the variance of the observed stimulation effects, a region of interest analysis was conducted. Using the bilateral stimulation as seeds, average functional connectivity to 12 regions of interest including the primary motor cortex, supplementary motor cortex, ventral anterior thalamic nucleus, parafascicular thalamic nucleus, globus pallidus externalis, globus pallidus internalis, substantia nigra pars reticulata, caudate nucleus, putamen, subthalamic nucelus, crus II cerebellum, dentate nucelus cerebellum^23–26^ was extracted for each patient. For each region, the average connectivity values were used to train a linear regression model predicting the stimulation effect. The estimated effect was then correlated with the empirical speed-dependent stimulation effect. P-values were corrected using the FDR-correction method^72^. Spearman’s correlation was used for the calculation of R-maps while Pearson’s correlation was used for quantification of explanatory power.

### Electrophysiology recordings

Bilateral subthalamic local field potentials (STN-LFP) using the remaining contacts and electrocorticography (ECoG) were recorded during the stimulation paradigm. Subdural ECoG strips were located contralaterally (n = 2) or ipsilaterally (n = 7) to the patient’s dominant hand which was chosen for the behavioral task. All data were amplified and digitized with a Saga64+ (Twente Medical Systems International; 4,000 Hz sampling rate) device. ECoG and STN-LFP signals were hardware-referenced to the lowermost contact of the left DBS electrode. Data was saved to disk for offline processing and converted to the Brain Imaging Data Structure for iEEG data, principally in brainvision format^73^. Data from patient 7, who performed the behavioral task contralaterally to the implanted ECoG strip, was utilized for exemplar analysis. Five ECoG channels located in proximity to the sensorimotor cortices as well as five contralateral LFP channels, were included in the analysis (see Figure 4). Custom Python scripts employing MNE-Python^74^, MNE-BIDS^75^, NumPy^76^, Pandas^77^ and Scipy^78^ were used for data processing and analysis (see Data and code availability section).

### Electrophysiological and behavioral data synchronization

Electrophysiology recorded with a SAGA64+ device and behavioral data recorded with MATLAB were synchronized using a custom pipeline in Python (see Data and code availability section). The time point of the first DBS artifact was visually identified and marked using the lowest available LFP channel in the right hemisphere. Then, the timepoint of stimulation onset in the behavioral data was identified and the mismatch between electrophysiological and behavioral data was subtracted from the behavioral data. The behavioral data recorded at ∼62 Hz was upsampled to the frequency of the electrophysiological recording by assigning the behavioral sample with the closest time stamp to each electrophysiological sample.

### Oscillatory power analysis

The signal recorded from the first ECoG electrode of patient 7, located over the motor cortex, was used to investigate changes in endogenous oscillations during STN-DBS. First, the signal was re-referenced by subtracting the average of all five ECoG electrodes (the sixth contact was excluded due to bad signal quality). Next, the data was epoched into 2-second windows centered on the time point of peak speed of each movement. Then, the data was transformed into the time-frequency spectrum using morlet wavelets with 4 cycles and normalized using z-scores with respect to the 400-100 ms preceding the peak speed. To ensure that differences in oscillatory power between stimulated and not-stimulated movements were not attributed to differences in kinematics, we identified a sample of not-stimulated movements that had the same size and did not differ in peak speed from the sample of stimulated movements. For stimulated trials, the DBS edge artifacts were replaced by NaN values using the time point of stimulation onset/offset for each movement and a standardized window of 65 ms. For visualization purposes, these NaN values were replaced with linearly interpolated values. Then, the average beta power (20-35 Hz) during stimulation and after stimulation was calculated for each movement (0.13-0.3 seconds, 0.41-1.0 seconds, windows based on the average onset/offset of the DBS artifact). The beta power during stimulated vs not-stimulated trials was statistically compared using a two-sided independent non-parametric permutation test with 100,000 permutations at an alpha level of 0.05. Further, significant differences irrespective of pre-defined time bins were examined using two-sided cluster statistics with 1000 permutations and an alpha level of 0.05. As this cluster analysis could not deal with NaN values and they were present during at least 1 epoch throughout the whole stimulation period, we replaced those values with the median of the remaining epochs if less than 15 % of epochs contained NaN Values, thus were affected by the DBS artifact. Time points during which more than 15 % of epochs were affected were excluded from the cluster analysis.

### Movement speed decoding

The movement speed of patient 7 during the behavioral task was decoded using five ECoG contacts located over the contralateral sensorimotor cortices and five LFP contacts placed in the contralateral subthalamic nucleus. ECoG signals were re-referenced by subtracting the average of all ECoG electrodes. LFP signals recorded from the second level of the DBS lead were summed together and bipolarly referenced against the lowest and highest contact resulting in two LFP channels (1-2/3/4, 8-2/3/4, with 1 being the lowest contact). The target variable was obtained by concatenating the raw speed values during each block. Decoding features were calculated using the py_neuromodulation toolbox in Python and included alpha (8-12 Hz), low beta (13-20 Hz), high beta (20-35 Hz), low gamma (60-80 Hz), high gamma (80-200) high-frequency activity (200-400 Hz) power computed by the Fast Fourier Transform and raw signal amplitude^30^. The frequency at which the features were calculated and the length of the segment used for calculation was optimized in a Bayesian optimization hyperparameter search to achieve maximum decoding performance (see Table S3). Further, the feature values were z-scored to the features in the preceding 1 s and clipped as an artifact rejection mechanism when they exceeded a value of [–3 3]. The machine learning algorithm CatBoost ^31^, which builds upon the theory of decision trees and gradient boosting, was used to predict the movement speed at each sample based on the instantaneous and a variable number of previous features. The number of boosting rounds was set to 30 and optimal tree depth, learning rate, and the number of preceding samples were identified using a Bayesian optimization hyperparameter search (see Table S3).

The decoding performance of 10 different channel combinations (1-5: individual ECoG channels, 5-7: individual LFP channels, 8: combined ECoG channels, 9: combined LFP channels, 10: combined ECoG and LFP channels) was tested using rigorous nested cross-validation. For each channel combination, the data was split into folds of 87.5 % training and 12.5 % test set using an outer 8-fold cross-validation. A Bayesian optimization search was conducted for 33 rounds using the training set only. For each optimization round, the training data from the outer loop was split into 75 % training and 25 % test set using an inner 4-fold cross-validation. The mean coefficient of determination R² averaged over all test sets of the inner loop was used to inform the hyperparameter search. Exemplar decoding features were visualized for an exemplar ECoG channel and movement. Feature importance, which measures how much the prediction changes when a feature is removed, was extracted from the CatBoost model trained on an exemplar outer fold with optimal parameters (the best model trained on all ECoG channels). The importance of preceding samples was averaged for each channel and feature.

## Data and code availability

Data can be made available conditionally to data sharing agreements following data privacy statements signed by the patients within the legal framework of the General Data Protection Regulation of the European Union. All code is made publicly available at: https://github.com/neuromodulation/speed_selective_DBS.git

