## Supplementary Information for "Differential modulation of movement speed with state-dependent deep brain stimulation in Parkinson’s disease"

### Supplementary Table 1: Subject details of 24 DBS patients and 14 healthy control subjects

| ***Healthy controls*** | | | **Parkinson's Disease patients** | | | | | | | | |
|---|---|---|---|---|---|---|---|---|---|---|---|
| ***n*** | ***Sex*** | ***Age*** | **n** | **Sex** | **Age** | **Disease Duration** | **Days After Surgery** | **UPDRS_III OFF** | **Dominant Hand** | **Dominant Disease Side** | **DBS parameters** |
| ***1*** | ***f*** | ***55*** | **1** | **m** | **57** | **6** | **4** | **26** | **R** | **R** | **R2-, 2 mA, L3-, 2 mA, 60 ms, 130 Hz** |
| ***2*** | ***m*** | ***61*** | **2** | **m** | **55** | **3** | **6** | **31** | **R** | **L** | **R2-, 1.5 mA, L2-, 1.5 mA, 60 ms, 130 Hz** |
| ***3*** | ***f*** | ***56*** | **3** | **m** | **45** | **5** | **3** | **14** | **R** | **Equal** | **R3-, 3 mA, L3-, 3 mA, 60 ms, 130 Hz** |
| ***4*** | ***f*** | ***68*** | **4** | **m** | **66** | **8** | **3** | **25** | **R** | **R** | **R3-, 2 mA, L3-, 2 mA, 60 ms, 130 Hz** |
| ***5*** | ***f*** | ***68*** | **5** | **f** | **65** | **6** | **3** | **41** | **R** | **L** | **R2-, 2.5 mA, L2-, 2.5 mA, 60 ms, 130 Hz** |
| ***6*** | ***f*** | ***62*** | **6** | **f** | **71** | **6** | **4** | **12** | **R** | **Equal** | **R3-, 2 mA, L3-, 2 mA, 60 ms, 130 Hz** |
| ***7*** | ***m*** | ***69*** | **7** | **m** | **54** | **12** | **6** | **28** | **L** | **R** | **R3-, 2 mA, L2-, 2 mA, 60 ms, 130 Hz** |
| ***8*** | ***f*** | ***48*** | **8** | **f** | **69** | **11** | **3** | **13** | **R** | **R** | **R3-, 2 mA, L3-, 2 mA, 60 ms, 130 Hz** |
| ***9*** | ***m*** | ***51*** | **9** | **m** | **53** | **9** | **3** | **27** | **R** | **R** | **R2-, 2.5 mA, L3-, 2.5 mA, 60 ms, 130 Hz** |
| ***10*** | ***f*** | ***64*** | **10** | **m** | **52** | **14** | **4** | **35** | **R** | **R** | **R2-, 2.5 mA, L2-, 2 mA, 60 ms, 130 Hz** |
| ***11*** | ***f*** | ***64*** | **11** | **f** | **73** | **5** | **4** | **32** | **R** | **R** | **R2-, 2 mA, L2-, 2.5 mA, 60 ms, 130 Hz** |
| ***12*** | ***m*** | ***53*** | **12** | **m** | **73** | **20** | **4** | **23** | **R** | **L** | **R2-, 2 mA, L3-, 1.5 mA, 60 ms, 130 Hz** |
| ***13*** | ***f*** | ***54*** | **13** | **m** | **50** | **10** | **3** | **31** | **R** | **L** | **R3-, 2 mA, L3-, 2 mA, 60 ms, 130 Hz** |
| ***14*** | ***m*** | ***60*** | **14** | **f** | **64** | **8** | **6** | **37** | **R** | **L** | **R2-, 2 mA, L2-, 3 mA, 60 ms, 130 Hz** |
| | | | **15** | **f** | **62** | **6** | **2** | **33** | **L** | **R** | **R3-, 2.5 mA, L2-, 2.5 mA, 60 ms, 130 Hz** |
| | | | **16** | **m** | **50** | **9** | **3** | **11** | **R** | **R** | **R2-, 2.5 mA, L2-, 2.5 mA, 60 ms, 130 Hz** |
| | | | **17** | **m** | **68** | **6** | **6** | **40** | **R** | **R** | **R3-, 2.5 mA, L3-, 2.5 mA, 60 ms, 130 Hz** |
| | | | **18** | **m** | **57** | **10** | **6** | **29** | **R** | **Equal** | **R2-, 3 mA, L2-, 2.5 mA, 60 ms, 130 Hz** |
| | | | **19** | **m** | **41** | **20** | **6** | **44** | **R** | **L** | **R3-, 2.5 mA, L2-, 2.5 mA, 60 ms, 130 Hz** |
| | | | **20** | **f** | **55** | **9** | **2** | **38** | **R** | **L** | **R2-, 3 mA, L2-, 2.5 mA, 60 ms, 130 Hz** |
| | | | **21** | **m** | **43** | **10** | **6** | **38** | **R** | **L** | **R3-, 2.5 mA, L2-, 2.5 mA, 60 ms, 130 Hz** |
| | | | **22** | **m** | **64** | **10** | **3** | **38** | **R** | **L** | **R3-, 2.5 mA, L2-, 3.5 mA, 60 ms, 130 Hz** |
| | | | **23** | **f** | **72** | **4** | **4** | **29** | **R** | **R** | **R2-, 2 mA, L2-, 3.5 mA, 60 ms, 130 Hz** |
| | | | **24** | **m** | **62** | **14** | **4** | **21** | **R** | **Equal** | **R2-, 3 mA, L3-, 3 mA, 60 ms, 130 Hz** |
| | ***9/15 F*** | ***59.50*** | | **8/24 F** | **60.50** | **8.79** | **4** | **26.79** | | **9/24 L** | |
| | | ***6.511*** | | | **8.95** | **4.28** | **1.13** | **8.57** | | | |

### Supplementary Table 2: Speed-classification accuracy

| | **Fast** | **Slow** |
|---|---|---|
| **Sensitivity** | 91.9 ± 7.0 | 95.6 ± 9.1 |
| **Specificity** | 98.4 ± 1.9 | 96.2 ± 3.8 |

### Supplementary Table 3: Bayesian Optimization Hyperparameters

| **Feature extraction hyperparameters** | |
|---|---|
| Sampling frequency | 20 to 50 Hz |
| Segment length | 200 to 500 ms |
| **CatBoost Hyperparameters** | |
| Number of preceding samples | 10 to 20 |
| Learning rate | 0.001 to 1 |
| Tree depth | 4 to 10 |

**Supplementary Figure 1: Comparison of average speed during stimulated and not-stimulated movement**

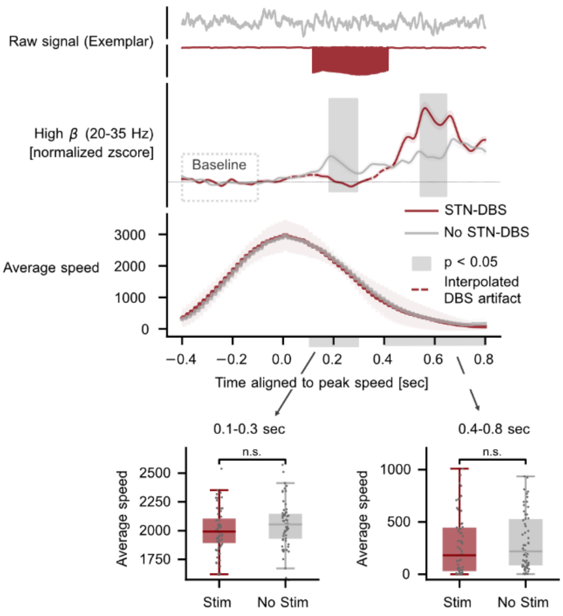

*Comparison of motor cortex beta activity during stimulated and not-stimulated movements of comparable peak speed demonstrates a stimulation-induced decrease in beta power during stimulated movements and a stimulation-induced increase in beta power after stimulated movements. Shaded light gray areas indicate clusters of significant differences. Shaded colored areas show the standard error of the mean. Differences between average movement speed of stimulated and not-stimulated trials during the peri-movement (0.1 – 0.3 s) and post-movement (0.4 – 0.8 s) period (gray bars below x-axis) are shown as box plots with trial-individual speed values. n.s. not significant*
